# R methylCIPHER: A Methylation Clock Investigational Package for Hypothesis-Driven Evaluation & Research

**DOI:** 10.1101/2022.07.13.499978

**Authors:** Kyra L. Thrush, Albert T. Higgins-Chen, Zuyun Liu, Morgan E. Levine

## Abstract

**Background:** Epigenetic clocks are promising tools for the study of aging in humans. The clocks quantify biological aging above and beyond chronological age, demonstrate systematic associations with risk factors that accelerate aging, and predict age-related morbidity and mortality. There is interest in using them as surrogate endpoints in intervention studies. However, the large number of clocks, decentralized publication and explosive popularity in the last decade has made for poor accessibility and standardization. This has hampered the abilities of new researchers to conduct truly hypothesis driven research—whether by not knowing about the best available clocks for a given question, or by systematically testing many or all as they become available.

**Results:** We report a centralized R package which can be installed and run locally on the user’s machine, and provides a standardized syntax for epigenetic clock calculation. The package includes a set of helper functions to assist with navigating clock literature and selecting clocks for analysis, as well as affording the user with the details of clock calculation. We describe each clock’s resilience to missing CpG information, combined with functionality to assess the need for imputation in the user’s own data. Furthermore, we demonstrate that while CpGs may not be shared among clocks with similar outputs, many clocks have highly correlated outputs.

**Conclusions:** Due to the previous decentralization of epigenetic clocks, gathering code and performing systematic analysis, particularly in protected datasets, has required significant information gathering effort. Here, we offer an R package with standardized implementation and potential for future growth and clock incorporation to assist with hypothesis driven investigation of aging as measured by epigenetic clocks. We show the potential of this package to drive the user to think globally about signals captured by epigenetic clocks, as well as to properly identify the potential and limitations of each clock in their current research.

## Background

Epigenetic clocks are promising tools, often discussed as future surrogate biomarkers for studies of aging and longevity. These clocks have been extensively reviewed; for their phenotypic associations [1, 2]; to understand the mechanisms of epigenetic aging [3, 4]; [5][6] Our current intent is to instead provide a practical overview of the categories, training methods, and applications of existing epigenetic clocks. As epigenetic clock research gathers further momentum in the study of aging, it is increasingly clear that a centralized toolkit to introduce the epigenetic clocks is essential. Such a toolkit must satisfy, in our estimation, a handful of requirements: It must (1) organize thematically and systematically the existing epigenetic clocks to minimize the risks of multiple testing and publication bias; (2) provide functionality to allow the researcher to perform not only *pro forma* analyses, implicating epigenetic clocks in a disease or dataset of interest, but push researchers to glean further biological insight as to the associations found; (3) be complete in its access of epigenetic clocks, while still giving editorial insight such as to make use of them navigable; (4) be sufficiently flexible so as to allow future advances to be made equally accessible.

Because of the relative ease of training epigenetic clocks and of DNA methylation (DNAm) collection, as well as the numerous age-related CpGs in the genome, there are currently numerous human epigenetic clocks available in the literature (Figure 1). The earliest such clocks utilize 1-10 highly age-associated CpGs in regression models, and these remain useful as low-cost assays [7–11]. However, the advent of large scale, streamlined collection of DNA methylation data on lllumina Beadchip methylation array technologies, as well as the adoption of elastic-net penalized regression to the training method, led to a new generation of clocks that can capture genome-wide aging signals. The first of these were trained to predict chronological age with high accuracy, including the Hannum blood [12] and Horvath multi-tissue [13] clocks, and have since expanded to include additional clocks [14–17].

**Figure 1:**
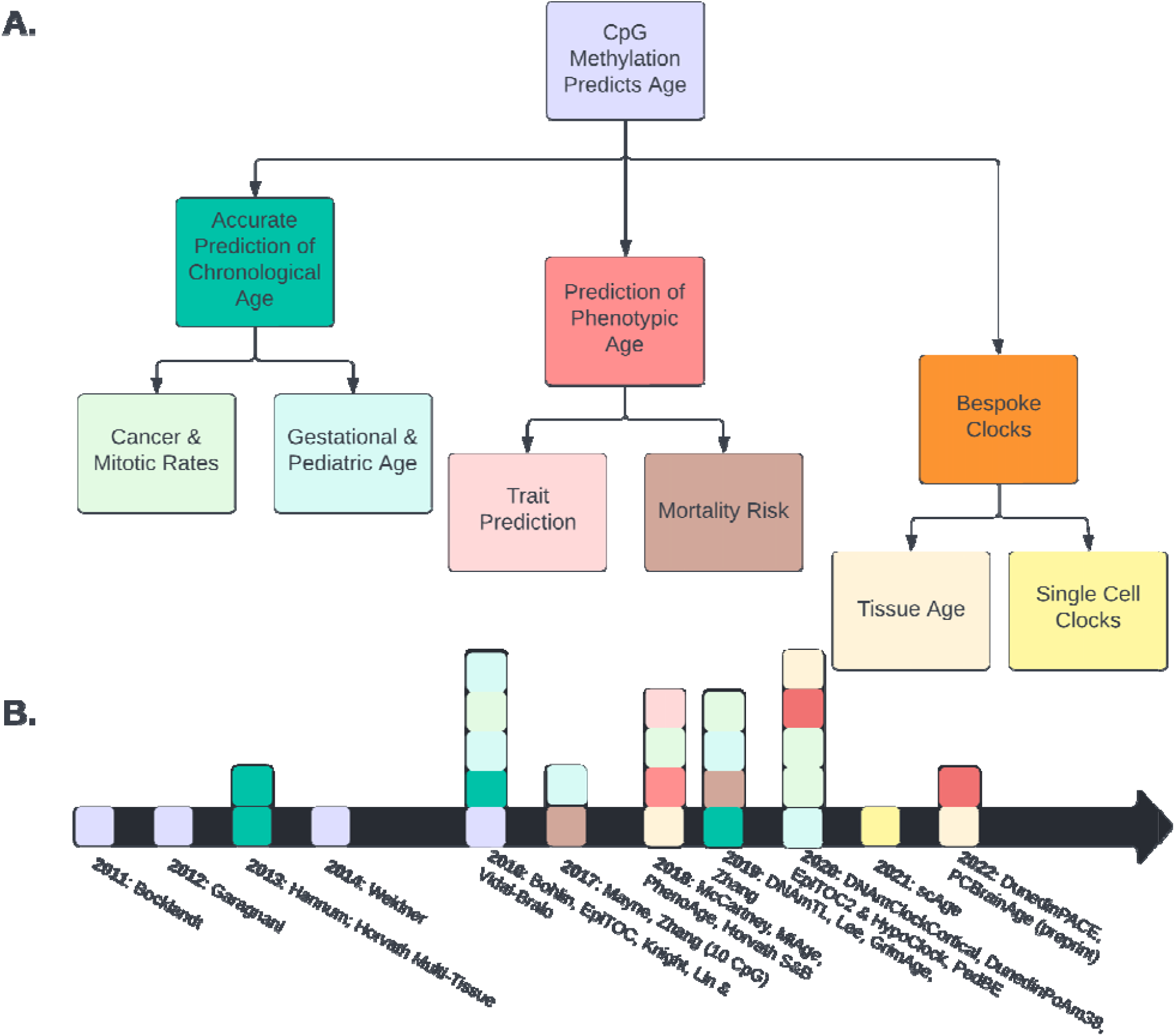
The Evolution and Diversity of Epigenetic Clocks. (**A**) Epigenetic clocks are semantically organized into key categories, and (**B**) individual clocks categorized and highlighted along a timeline. Note that years with multiple clocks are colored in blocks from top to bottom according to the alphabetical list.

For those studying development and gestation, significant effort has been spent to create reliable gestational and pediatric age clocks [18–22]. Similar approaches led to the generation of mitotic clocks, so-called for their presumed ability to track the rate of mitotic divisions and project cancer risk [23–25]. The telomere length estimator DNAmTL, which is highly correlated with cellular replication rate, can be included in this category as well [26].

In addition to clocks that predict discretely measurable aspects of age and cell turnover, efforts have been made to capture heterogeneity in aging as meaningful biological signal. These [27][28], and DunedinPACE[29] are trained to predict individuals; degree or rate of biological change with time, especially those changes that contribute to age-related morbidity and mortality risk. It has also been found that DNAm can be used to predict various traits, lifestyles or exposures that are not necessarily related to aging [30].

Finally, there has been recent interest in “bespoke” clocks designed for particular tissues, diseases, data types, or applications. For example, one ongoing challenge for epigenetic clocks is that most human clocks were trained using primarily whole blood[2]. Multi-tissue clocks[13]s conserved across tissues and may ignore tissue-specific aging changes [6]. Thus, tissue-specific, bespoke clocks have been developed, including for skin-and-blood [16], brain cortex [31], skin [32], and the scAge framework for predicting epigenetic age from single cell methylation [33] have been reported. Also, since clocks are often trained in large aging cohorts, it is possible they may miss aging patterms that occur in small subsets of the population, as in the context of rare diseases. Due to the increasing abundance of non-blood tissue DNAm, new methods for collection [34], additional approaches to clock-training [35, 36], and emerging age-related diseases, we predict that the number of bespoke clocks will see a dramatic increase in the next few years.

Yet the challenge remains: Which clock should be used? The variety of epigenetic clocks can be useful for investigating many different aspects of aging. But selecting the appropriate clock(s) for a study requires navigating a decentralized body of nuanced literature. The choice of clock may be impacted by the phenotype that they were trained to predict or the context they were trained for. However, the differences between clocks can be subtle, amounting to differences in training data composition or procedure, such as age ranges [22], or preselection of CpGs [23].

This clock selection problem creates concerns regarding the integrity and interpretability of aging studies. In particular, there are two consequences we would hope to avoid. The first is exclusive, repeated testing of the best cited and most reported aging clocks—namely the Horvath multi-tissue, Hannum, Levine PhenoAge and GrimAge DNA methylation clocks. [1]. This aligns well with the plethora of publications reporting the associations of acceleration of the Horvath multi-issue, PhenoAge, and GrimAge predictors with biological changes[37–39], and disease risk or mortality[40–45]. While this produces some standardization in the field, these clocks are not necessarily the optimal choice in all cases. If a researcher instead has all available clocks at their disposal and then applies a hypothesis-driven selection of clocks, an alternative, lesser-known clock may indeed be the optimal choice. The second unintended consequence could be that individuals test many clocks as clocks are published or the researcher becomes aware of them, and only significant results tend to be noted and published.

The variety of clocks and their decentralized distribution also creates practical obstacles for aging research. Researchers wishing to apply epigenetic clocks must first mine the literature for their options, identify one (or multiple) clocks to test, locate and download the data to do so, and ensure that the calculation is properly performed across their samples. This process creates substantial logistical barriers for researchers. Clocks that are published along with public code and data should be applauded. Furthermore, a few existing platforms can calculate multiple clocks. These include the online Horvath calculator (http://dnamage.genetics.ucla.edu) and EstimAge (https://estimage.iac.rm.cnr.it). However, these require data to be uploaded to a third party server, which is prohibited for protected datasets and limits researchers’ access to the underlying details of calculation. There also exists the methylclock Bioconductor package, which is currently limited to chronological age clocks, gestational age clocks, DNAmTL and PhenoAge. In summary, the sheer number and variety of clocks creates two primary challenges that impede use by the broader scientific community: (1) the selection of the most appropriate clock(s) for the scientific question or hypothesis at hand; (2) access to the many clocks. We address this by providing a centralized resource in which individuals can explore, investigate, and calculate any and all clock(s) appropriate for a research question from a project’s inception. This is a necessary improvement for the field, as it allows for systematic study of epigenetic clocks, which in turn advances future understanding of their underlying relationships and biological significance.

There is currently no true standard format or resource for the researcher to publish and distribute clocks. Here we present a consolidated resource that applies a standardized format to the calculation of epigenetic clocks, establishes a repository for the fitted values of existing clocks, and provides a few helper functions for the exploration of appropriate clocks, their CpGs, and inter-clock correlations. Further, this package can be installed and run on a local machine, eliminating the need for uploading of potentially protected data, and affords near-immediate results to the researcher, regardless of the number or identity of clocks they choose to select. To further facilitate accessibility of the epigenetic clocks, we have also provided a thorough tutorial walking through use of the package and questions to be addressed on the github page for this package (github.com/MorganLevineLab/methylCIPHER).

## Implementation

*This should include a description of the overall architecture of the software implementation, along with details of any critical issues and how they were addressed*.

The current package is implemented using the R programming language and distributed via installation from Github (github.com/MorganLevineLab/methylCIPHER). This distribution allows us to provide a flexible, regularly updated, and community driven package. Not only can we push regular updates to users as new clocks are added, but the research community can rapidly suggest new clocks, helper functions, or improvements to code. Users wishing to generate their own independent Github R packages during the publication process of novel clocks can be imported by this package, or be referred to in the online Github based wiki and tutorial. The functions of this package are represented in the schema in Figure 2.

**Figure 2:**
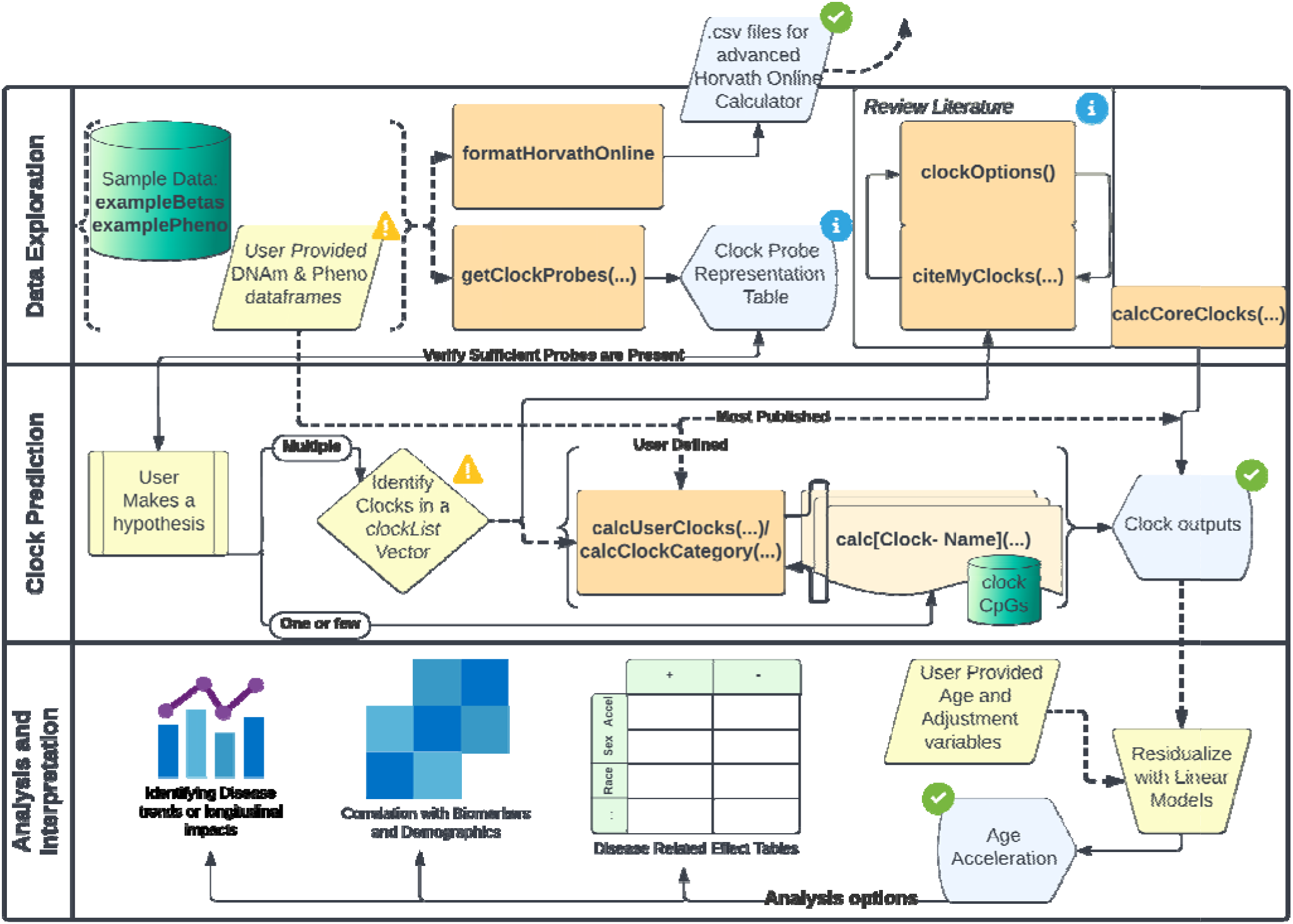
methylCIPHER Function Schema. A representational schematic of the functionality contained within methylCIPHER. Data is connected as inputs with dashed lines. User inputs are colored in yellow, with objects required to be supplied by the user highlighted with “!”. Orange rectangles indicate functions exported by methylCIPHER for the user to run. Blue pentagonal boxes indicate outputs, with green checkmarks as endpoints. Green cylindrical objects are RData objects stored and accessible from within the package.

To calculate epigenetic clocks in methylCIPHER (functions of the form *calc[<u>Clock-Name]</u>*), the user provides a labeled data matrix of pre-processed methylation Beta values obtained via the Illumina HumanMethylation450 Beadchip or Infinium MethylationEPIC kit from Illumina (San Diego, CA). These are commonly referred to as 450k and EPIC arrays respectively. Preprocessing and normalization of methylation data is typically performed within R using the *minfi* [46], *wateRmelon* [47], or *SeSAMe* [48] packages, however methylCIPHER functions regardless of normalization protocol. For more details regarding the effects of choice of normalization, refer to Ori et al. [49]. The user must simply have an object of *matrix* or *data frame* class, with named columns corresponding to the Illumina CpG names, and cells containing methylation beta ratios between 0 and 1.

We recommend that the methylation data object have named rows corresponding to unique sample identifiers. Most analyses will benefit from a corresponding “phenotype” data frame with sample identifiers; sample metadata; demographics; health outcomes; age; sex; and other traits of interest. While optional for individual clock calculations, this typically assists the researcher with downstream analyses. Without this data frame, clocks can be computed using only a single function at a time, with output to a vector object. However, the “pheno” dataframe provides a central location to append multiple clocks to if using the multi-clock wrapper functions *calcUserClocks* or *calcCoreClocks*.

Of note, for some analyses (for example, calculation of IEAA or EEAA [50]) estimates of blood cell composition are necessary. To obtain such estimates, individuals may want to use the Houseman method [51] of cell-type deconvolution from the *minfi* package. However, local methods for predicting cell composition of blood can be effectively run only when preprocessing occurs from raw methylation files (i.e. IDAT). If access is limited to preprocessed methylation beta values, as in some publicly available datasets, the Horvath Online calculator can predict blood component proportions. We hope to provide users with this convenient functionality in the future. However, we have provided a formatting function *formatHorvathOnline* which allows the user to quickly generate the input files for the Horvath Online calculator. This allows for both calculation of blood composition estimates and GrimAge [28].

The current version of the *R/methylCIPHER* package has been tested on both Windows and Mac computer systems, running R v3.6+. It can run on most modern personal computers, requiring less than 16 GB of RAM (and in many cases less than 8 GB) for methylation datasets containing hundreds of samples at once. The functions run efficiently and will provide near-immediate results within seconds or minutes.

## Results & Discussion

The R package *methylCIPHER* provides both seasoned and casual users of epigenetic clocks with the tools necessary for thorough, hypothesis-driven research using existing epigenetic clocks. We provide a comprehensive listing of human epigenetic clocks that use; (1) linear approaches; and (2) CpGs found in the commonly utilized Illumina 450k and EPIC arrays. This broad set of clocks can be searched through the function *getClockOptions()*, which allows users to explore their options. We have also provided convenient referencing of the source papers for each of the clock calculation functions, using *citeMyClocks*. This accepts a group or list of functions at once, which in turn allows readers to quickly refer to the original clock papers and understand the underlying principles of their training. This process of information gathering is shown graphically in the top right region of Figure 2.

Due to the variable performance of experimental designs, and a multitude of existing pipelines for quality control, users may find missing probes or DNAm values in their data. This can be seen as missing probe columns in the final normalized beta value matrix, columns of *NA* values, or sporadic *NA* values in sample/ probe pairs. If the probes are missing entirely from the beta value matrix, this can impact the decision to implement specific clocks. Therefore, *getClockProbes* provides the user with a table to determine what portion of probes are available for the various clock options, so that a clearly informed decision can be made. Alternatively, they may find columns of all NA values, which can be removed using *removeNAcol*. Sporadic missing values for select probe/sample pairs can either be mean imputed across the matrix samples (using function *meanimpute*), or imputation can be done later within the clock calculation functions if a mean vector is provided as a reference containing the necessary CpGs. However, most clocks can be calculated without imputation by simply ignoring those CpGs in the resultant weighted regression values for such samples. The effect of doing so varies by clock.

In figure 3 we visualize the degree of information contained in individual CpGs. The model contribution of each CpG was estimated by multiplying the absolute regression value in each clock for each site by its standard deviation in the Framingham Heart Study (FHS) offspring cohort [52], and plotted against the standard deviation alone. These results are shown for the Hannum (Figure 3A), Horvath Multi-Tissue (Figure 3B), and PhenoAge (Figure 3C) clocks (additional clocks in supplemental materials, figures S1-S2). With the CpGs plotted in this manner, we can see that each CpG in the Hannum clock tends to have higher weight, evident from the higher mean contribution represented by the blue horizontal dashed line. Further, PhenoAge employs CpGs with higher standard deviation than the other clocks, but due to lower weight in the clock regression, these tend to have dampened model contributions. These plots help to conceptualize the effect of using mean imputation on model CpGs, as utilizing mean imputation removes signal in individual samples that may reflect meaningful inter-individual differences.

**Figure 3:**
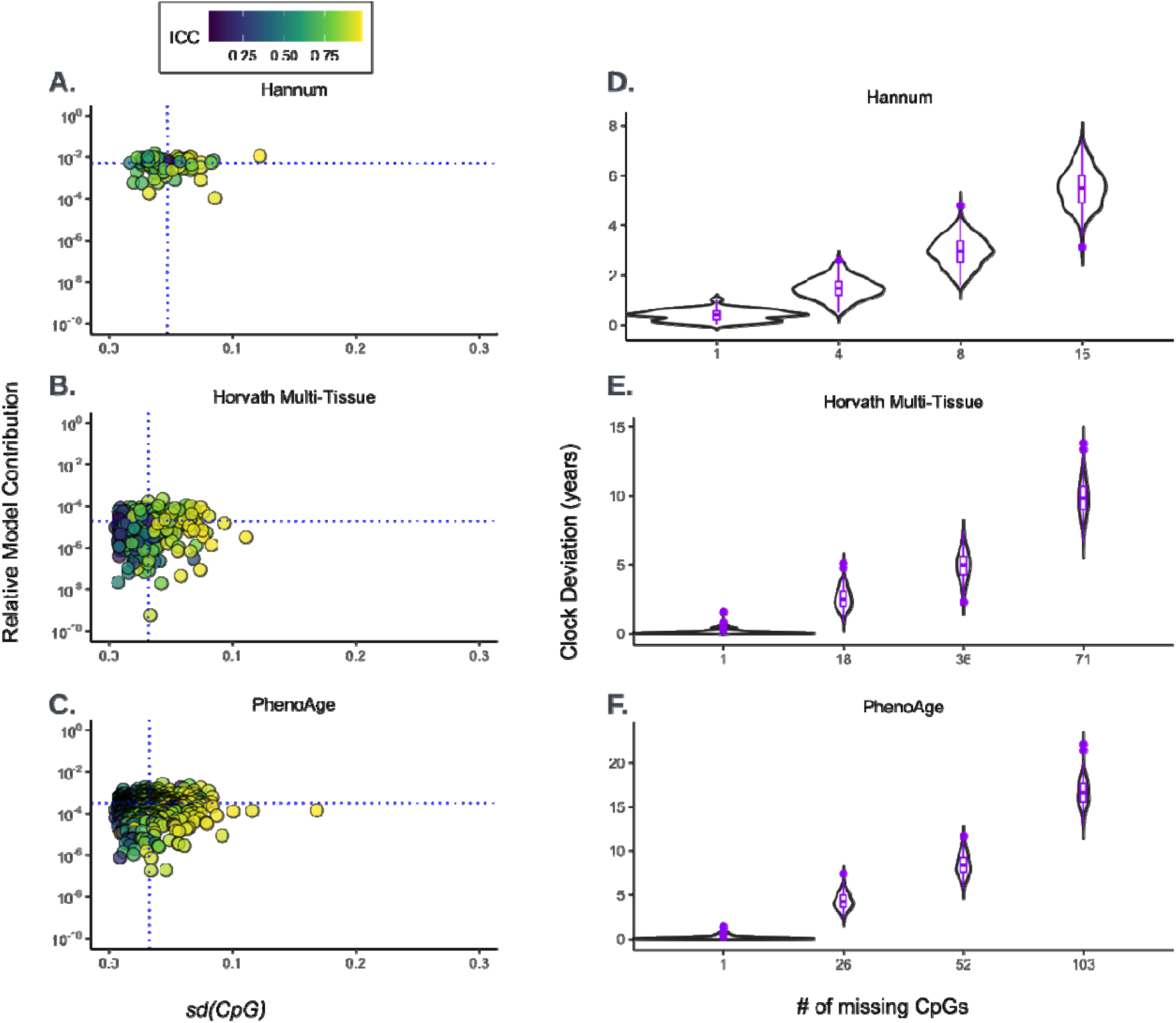
Impacts of Missing CpG Imputation on Key Clocks. Three representative clocks, Hannum (**A, D**), Horvath Multi-Tissue (**B, E**), and PhenoAge (**C, F**), were selected to assess the effects of imputation. Relative model contribution was calculated according to the absolute CpG weight in the regression model, multiplied by the standard deviation of the CpG in the Framingham Heart Study dataset. The model contributions of each CpG were plotted against the standard deviation of the CpGs, with the mean of each axis plotted as a blue crosshatch (**A**-**C**). Imputation of CpGs in the top right quadrant, with high standard deviation and high model contribution, will have a greater impact than CpGs in the other quadrants. Further, clocks whose CpGs extend further into this region will be more impacted by mean imputation effects. We repeatedly tested and plotted the effects of mean imputation on 0.1%, 5%, 10% and 20% of CpGs selected at random (**D**-**F**). Due to varying sizes of the clocks, these percentages represent varying numbers of missing CpGs for each of the clocks.

To functionalize this effect, we simulated the effects of increasing amounts of missing array methylation probes for a given sample. We performed 1000 iterations each of randomly drawing 0.01, 5, 10, and 20% of CpGs within each clock. Then, we found the distribution of *Clock Years* information contained within each potentially missing sample. Because we are approximating information lost, the *Clock Years* measurement is defined by the absolute value of the sampled CpGs’ clock regression coefficient, multiplied by the standard deviation of these CpGs. The result is an approximation of the information in years of measurement lost for a sample or set of samples for which that proportion of CpGs’s methylation is unavailable. We note that because *Clock Years* is defined as absolute values for each CpG, oppositely weighted CpGs in a clock do not counteract each other. Thus, the information lost may reflect a larger difference than the shift in actual clock value. As each clock selected is of varying size, the percentage of missing CpGs varies in the absolute number lost. Here, we demonstrate that *Clock Years* information lost appears strongly associated with clock size: Bigger clocks show a larger amount of information lost, despite removal of the same proportion of sites (Figure 3D-F). However, as is demonstrated in the disparate axis scaling between clocks, there must be far more CpGs missing to exert a similar effect on the larger clocks than Hannum.

The user can choose, based on what is most appropriate for their hypothesis and data available, a set of CpG-based DNA methylation clocks to calculate. Then, either using the manual functions of the form *calc[<u>Clock-Name]</u>*, or a user-specific character vector list of clocks, input to *calcUserClocks*, the appropriate values can be generated. These are ideally output by binding to an existing “phenotype” data frame supplied by the user containing relevant sample metadata.

To calculate each clock, an RData object is accessed containing CpG identity and weight information as supplied by the original authors. Each of these objects can be accessed using *data(“[<u>Clock-Name]</u>_CpG(s)”)*. A central repository where these data objects are readily accessible has two advantages. First, it makes the details of clock calculation transparent to the user. Second, it facilitates studies of clock CpG identities and their biological underpinnings.

For example, methylCIPHER allowed us to quantify the overlapping CpG identities of clocks within some of the categories identified in Figure 1. It is often discussed within the field that CpGs—at least those on the Illumina array technologies— change in a concerted, multi-collinear manner with age. This has motivated some of our prior approaches to clustering clock CpGs to ascertain underlying biological signals or changes [53, 54]. We find that the vast majority of CpGs selected by clocks do tend to be unique to those clocks, though a small subset of methylation sites are common across many clocks within categories (Figure 4A-B). Other observatons, such as the fact that EpiTOC2 is a subset of the original EpiTOC sites, or that DNAmTL is entirely unique in its CpG selections, are immediately obvious from this analysis (Figure 4B). However, despite sparse overlap in clock CpGs, it remains that these clocks’ sex-adjusted age accelerations (i.e., residuals of regressing clock values onto sample age and sex) are typically quite correlated (Figure 4C).

**Figure 4:**
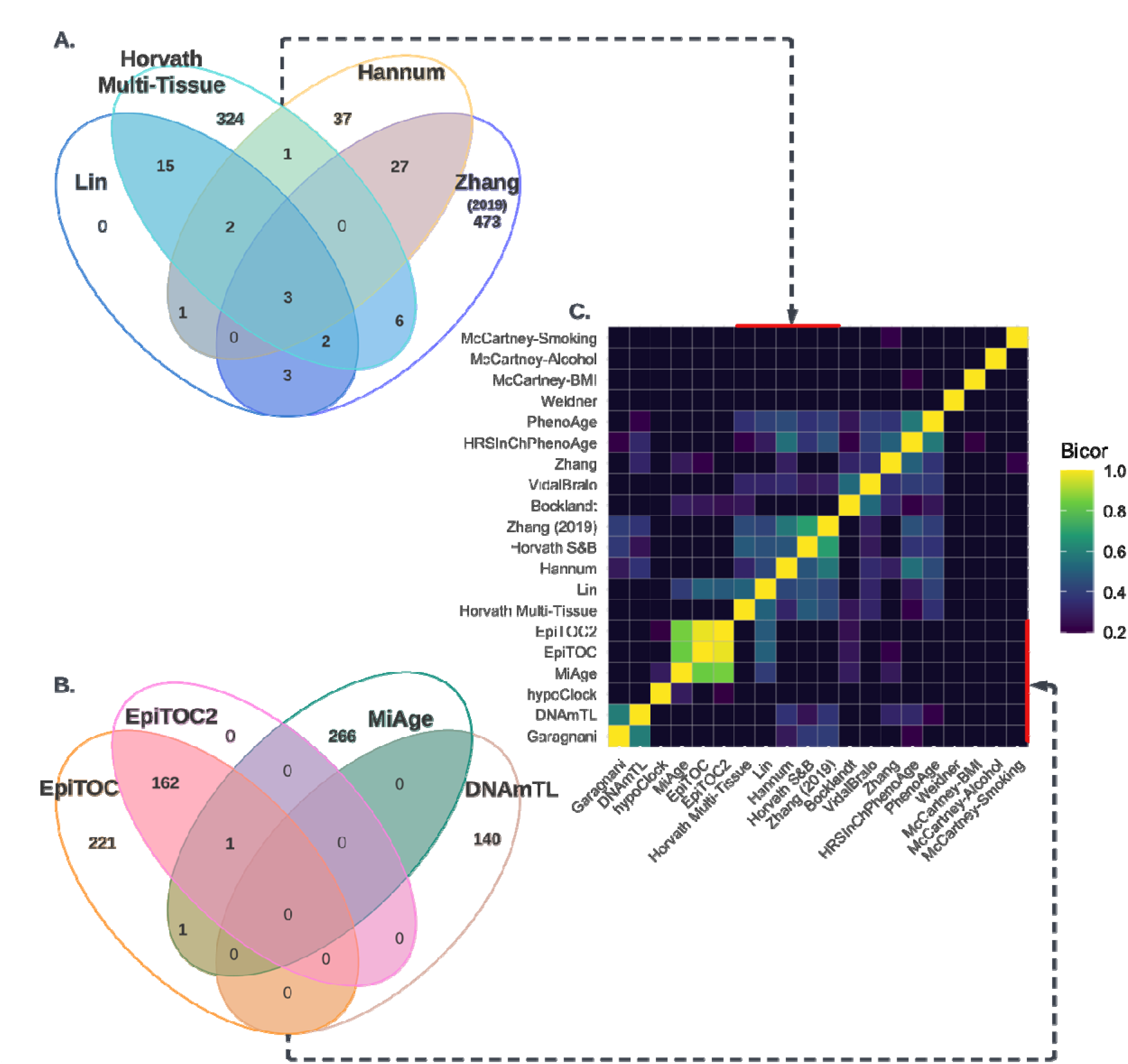
Shared Signal Does Not Arise From Shared CpGs. Epigenetic clocks were selected from the categories of highly accurate chronological age clocks (A) and cancer & mitotic rate clocks (B) as defined in Figure 1. Selection of CpG identity overlap was limited to 4 clocks per category, and in the case of mitotic clocks hypoClock was not included as it was designed to select different CpGs from EpiTOC2. (C) Sex-adjusted clock residuals were found in FHS and correlated according to biweight midcorrelation (thresholded at >0.2). While clock residuals can capture well-correlated information, their CpG overlaps within a cluster can be quite low.

In fact, it was recently reported that numerous combinations of CpGs can be used to train epigenetic clocks across the epigenome, a concept which arises primarily from this noted redundancy [55]. To further investigate the CpG identities of the clocks as distributed by the original authors in our data files, we repetitively retrained multiple well known epigenetic clocks. We performed this analysis in both a chronological aging clock (Hannum) and a tissue specific aging clock (Horvath Skin & Blood). We first generated an experimental design matrix (Figure 5A), which consists of 19×200 sample cells. Each cell is the result of one of 19 bootstrapped samples of the training data individuals drawn with replacement (for up to 6 draws of the same individual), and 1 of 200 versions of 10,000 CpGs drawn without replacement from the available probes on the 450K array. The concept of bootstrapped samples for model training was inspired by the original Hannum training method [12]. We used the original Hannum training dataset [12], and the publicly available training datasets (supplemental materials, table 1A) from the original Horvath Skin & Blood clock publication.

**Figure 5:**
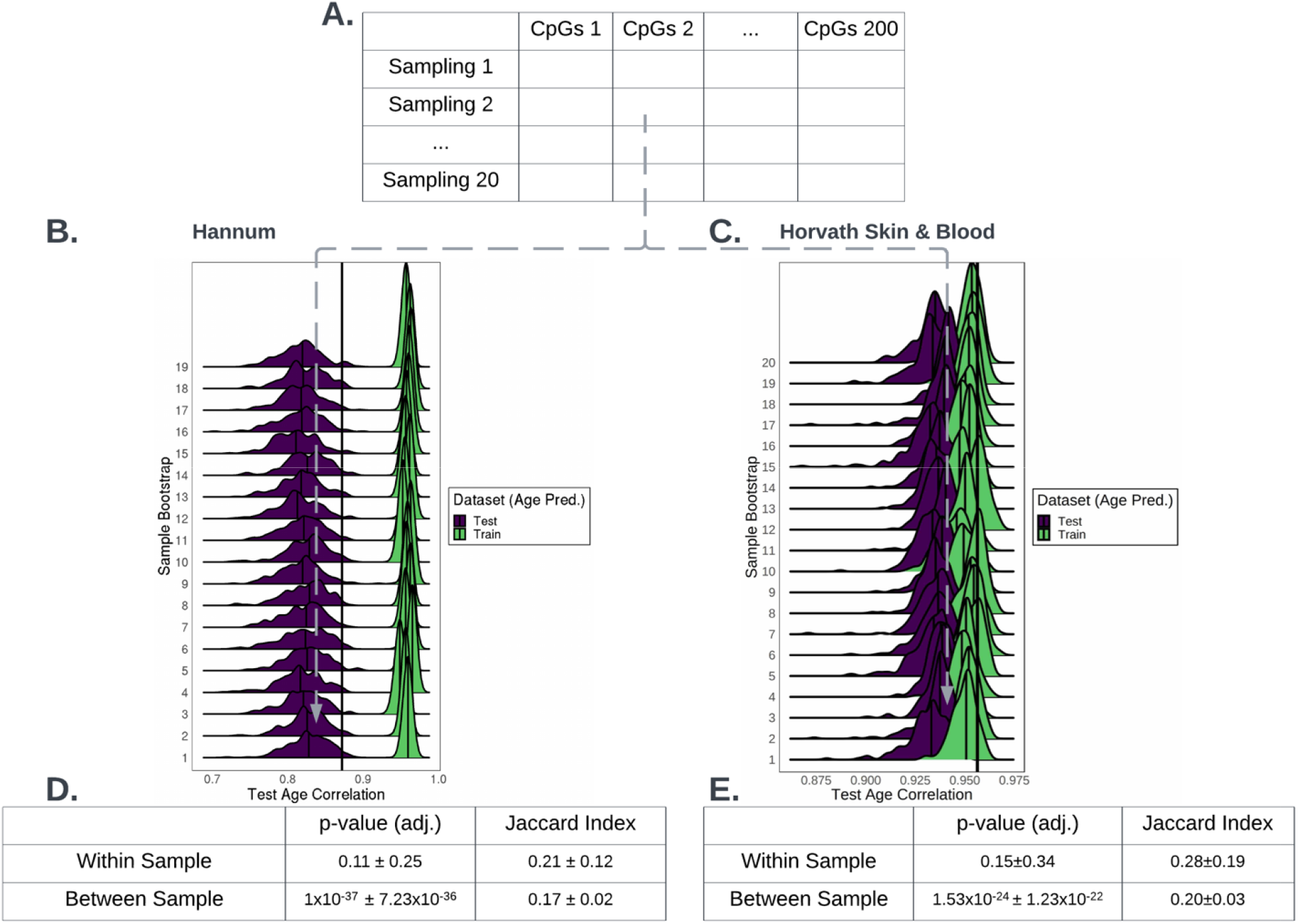
Bootstrapping Methylation Data Demonstrates Methylation Redundancy. The experimental design matrix (**A**) demonstrates how both samples (19x) and CpGs (200 × 10k CpGs) were selected with random amounts of overlap in their sampled dimensions. Each sampling cell was used to retrain an elastic net regression model of the Hannum (**B**) or the Horvath Skin & Blood (**C**) clocks. The large vertical lines demonstrate the correlation of the originally developed clock in the whole test dataset. Smaller vertical lines in the purple density plots indicate the median age correlation of the within-sample bootstrap elastic net trials. As each experiment is allowed to overlap in CpGs to an extent, we use the p-value and Jaccard indices of modified gene set overlap tests to determine whether CpGs are repeatedly selected across bootstrapped CpG lists and within a given list across sampling sets, for both Hannum (**D**) and Horvath Skin & Blood (**E**).

Given each experimental matrix cell, we retrain the epigenetic clocks in the sampled data, by applying elastic net penalized regression on chronological age. Elastic net regression was performed with 10-fold cross validation and a 0.5 ratio of LASSO and Ridge regression. All models were subsequently evaluated in an independent test dataset consisting of either whole blood methylation data [56] or skin and fibroblast datasets (supplemental materials, table 1B).

This analysis enabled comparison across randomly selected subsets of individuals and/or CpGs on retrained clocks. The correlations of predicted age and true age measures were visualized as density plots organized by sample bootstrap, for both Hannum (Figure 5B) and Horvath Skin & Blood (Figure 5C). Training sample correlations were verified normal distributions with high degree of correlation, while independent test sample correlations for these models retained relatively high correlations. Medians of each density plot (small vertical bars) are compared to the original clock’s correlation to age in the test dataset (large vertical bar). As is demonstrated in the case of Hannum, while some CpG subsets produce models that have correlation as low as 0.89 with chronological age, this is still a relatively high correlation. Thus, while access to some CpGs improves model performance, the improvement is modest. Furthermore, even when lists of CpGs are partially overlapped, the degree of overlap in selected CpGs is lower than expected (Figure 3D). Therefore, while some CpGs may contain relatively important information for age prediction, there may be significant redundancy in DNA methylation, allowing high model correlation and performance even with low shared identity of CpGs. It is also important to note that the original Hannum methylation age clock used CpG preselection [12] whereas here we are performing unsupervised selection. This may account for the overall slightly reduced performance. However, we find it more promising that the spread of resultant correlations in test data is relatively small, retaining correlations above 0.8 for the vast majority of models. This high redundancy amongst CpGs may largely explain why different clocks trained to predict the same outcome can have such sparse overlap in their composition.

Similar results are found for a tissue specific aging clock, Horvath Skin & Blood (Figure 3C,E). Here, we see that despite training and testing with half the sample size of the original clock, more than 50% of the models outperform the original clock’s prediction in the test dataset. It has been demonstrated that there are measurable differences in age related DNA methylation changes between tissues [57, 58]. Consequently, there may exist particular sets of CpGs which are essential to the function of for tissue specific aging clocks [49]. However, our results suggest that the majority of the CpGs have significant redundancy even for use in a tissue specific age predictor. Again, the selected CpGs do not have significant overlap between lists, despite them having significant overlap within a list across sample bootstraps (Figure 5E). Therefore, there may be many CpGs to select from for age prediction in tissue-specific contexts, though given the limitations of the current training method, they may not be truly tissue-specific merely single-tissue trained.

Beyond enabling characterization and investigation of the existing epigenetic clocks’ mechanics, the current package enables efficient comparison of desired clocks with important biological phenotypes, biomarker data, or other sample metadata (Figure 6). Due to the rapid calculation of DNAm-based clocks in new data with automatic appending of results to existing phenotype/ sample metadata, researchers can quickly search for associations between outcomes or biomarkers, and clock scores.

**Figure 6:**
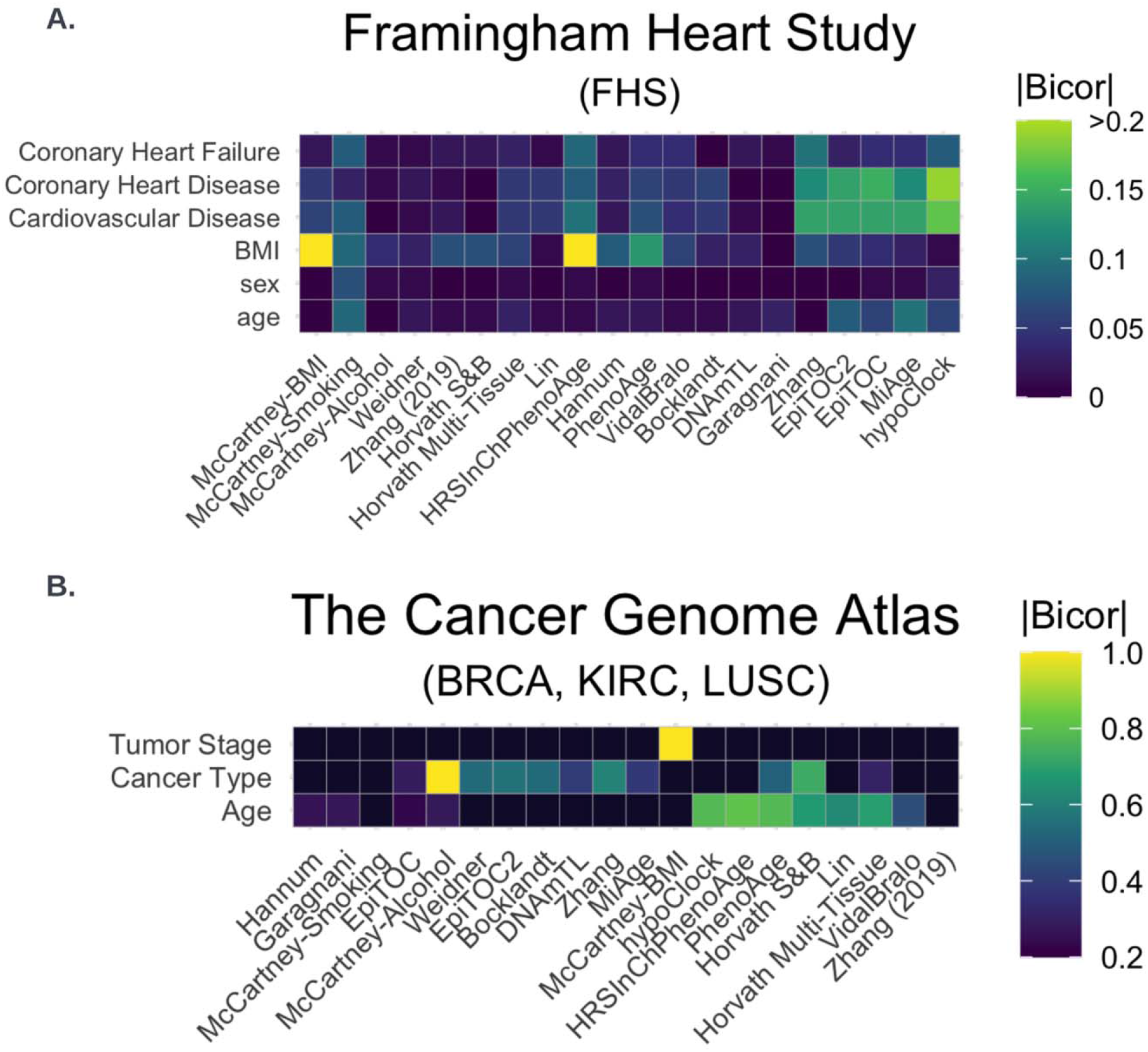
Clock Univariate Associations Capture Key Signal In Diverse Datasets. The available epigenetic clocks’ simple age regression (regression of clock onto just age) were converted to z-scores across each dataset, as were samples metadata such as age, BMI, or sex. Additional sample traits were left as original variables as z-scores aren’t realistic. The univariate associations were then described as the absolute biweight midcorrelation between the z-scored clock residuals and sample metadata. (A) In FHS data, there is clear associations between traits of interest and a cluster of mitotic clocks, whereas (B in a few cancers in TCGA data, McCartney lifestyle/ trait predictors of Alcohol and BMI show the strongest correlations to traits of interest.

Typically, the user will want to residualize the clock scores rather than using raw scores, to assess the effects of age acceleration. We have left this step up to the user, as details of correction for batch, sex, race, or other features tend to be dataset, and use specific. For instructions on how to approach calculating age residuals, please refer to the included tutorial or wiki on our GitHub distribution repository.

We find that we can rapidly uncover interesting results, such as univariate associations between the cluster of accelerated mitotic clocks and cardiovascular outcomes in the Framingham heart study (FHS) data (Figure 6A). Alternatively, we find that the acceleration of BMI and Alcohol clocks by McCartney et al. [30] well surpass the univariate associations found by other clock accelerations to cancer types and stages in a subset of TCGA data (Figure 6B).

In nearly all cases, we do not recommend that the user calculate all clocks available to them, as this introduces significant multiple testing. We have previously shown that clocks can be driven by similar information content [53], highlighting the potential utility of clustered-clock approaches. Typically the researcher would be best served by selecting a hypothesis-driven subset of clocks: This would look like a single clock tested from within a cluster (e.g. one mitotic clock) or calculating and reporting all clocks and their agreement. However, we aim to show the ease with which few or all clocks can be assessed using the *methylCIPHER* package, as well as to provide resources to guide those decisions according to the high degree of correlation between clocks, particularly relative to those of the traits of interest.

## Conclusions

The current software is an important compendium of clocks currently distributed through a wide variety of means. This reduces impediments to users, both in gathering the data for calculation, and ensuring reproducible and accurate calculation. Further, prior decentralized reporting and distribution of epigenetic clocks has led to the potential for researchers to inadvertently conduct significant multiple testing, potentially without proper correction: This can even occur throughout the course of a project in which the researcher becomes aware of, and iteratively tests, additional epigenetic clocks.

Through the provision of standardized clock calculation functions, and tools to rapidly investigate options available to the user, we aim to improve uptake of epigenetic clocks while enhancing the reproducibility of DNAm clock-based studies in the future. Further, as the current package is installed locally to one’s personal computer or computing cluster, it is possible to rapidly calculate several epigenetic clocks, even in protected data. We intend to expand the clocks contained in this package in the future: (1) The addition of future human DNAm (regression) based clocks to the present researcher’s toolkit will be essential; (2) Availability of mammalian arrays [34] will spur the use of similarly implemented epigenetic clocks for nonhuman vertebrates, and should also be included here; (3) The online tutorial provided in the currently discussed Github repository will be further expanded according to developing practice and user suggestions for standard features of epigenetic clock-based disease and trait analysis. Due to their different operating requirements and less standardized implementation, other forthcoming methods of epigenetic clock calculation are unlikely to be housed within this package, but we will direct users to their own sources using our wiki and tutorial pages. These include, but are not limited to, deep learning-based clock approaches [35, 59], the next generation of low-noise clocks referred to as PC Clocks [36], and single cell epigenetic clocks approaches [33].

## Availability and Requirements

Project name: methylCIPHER

Project home page: github.com/MorganLevineLab/methylCIPHER

Operating system(s): tested on Mac OS 11+, with and without M1 chip, Windows

Programming Language: R 3.6.1+

Other requirements:

License: TBD

Any restrictions to use by non-academics: TBD

## List of abbreviations

## Declarations

### Ethics approval and consent to participate

Not applicable

### Consent for publication

Not applicable

### Availability of data and materials

All data necessary for clock calculation is provided through the distributed package. Datasets used for figures 4 and 6 were provided by the Framingham Heart Study (FHS) Offspring Study Cohort and The Cancer Genome Atlas (TCGA). The data used for Figures 4C and 6A in the current publication are based on the use of study data downloaded from the dbGaP web site, under phs000724. TCGA data (Figure 6B) can be downloaded via the GDC Data Portal (https://portal.gdc.cancer.gov).

### Funding

#### Authors’ contributions

KLT was responsible for conception, packaging of code, and writing the present manuscript. AHC, ZL, and MEL contributed code and suggestions for clocks to include within the package, and reviewed the manuscript.

## Acknowledgements

All figures were created in whole or in part using Lucidchart.

