## Supplemental Materials for "R methylCIPHER: A Methylation Clock Investigational Package for Hypothesis-Driven Evaluation & Research"

| S1A. Hannum Retraining Data | | | | | |
| --- | --- | --- | --- | --- | --- |
| Dataset | Reference | Tissue | Use | N | Age Range |
| GSE40279 | Hannum 2013^1^ | Blood WB | Training | 656 | 19-101 |
| GSE41169 | Horvath 2013^2^ | Blood WB | Test | 94 | 18-65 |
| S1B. Horvath Skin & Blood Retraining Data | | | | | |
| Dataset | Reference | Tissue | Use | N | Age Range |
| GSE77136 | Ivanov 2016^3^ | Dermal fibroblasts | Training | 43 | 0.1-85 |
| GSE52025 | Wagner 2014^4^ | Fibroblasts | Training | 62 | 23-63 |
| GSE79056 | Nashville Birth Cohort | Blood Cord | Training | 36 | -0.3-0.02 |
| GSE104471 | Yang 2017^5^ | Epithelium and PBMC | Training | 72 | 24-74 |
| GSE50759 | Berko 2014^6^ | Buccal | Training | 96 | 1-28 |
| GSE80261 | Portales-Casamar 2016^7^ | Buccal | Training | 215 | 5-18 |
| GSE51954 | Vandiver 2015^8^ | Epidermis or Dermis | Test | 37 | 20-84 |
| GSE62924 | Rojas 2015^9^ | Blood Cord | Test | 38 | -.1-0.04 |
| GSE80283 | Victorian Infant Collaborative Study | Blood Cord | Test | 183 | -0.3--0.1 |
| GSE94876 | Jessen 2019^10^ | Buccal | Test | 120 | 35-60 |
| GSE109042 | Lussier 2018^11^ | Buccal | Test | 51 | 3.5-18 |
| GSE111223 | Chuang 2017^12^ | Saliva | Test | 259 | 36-88 |

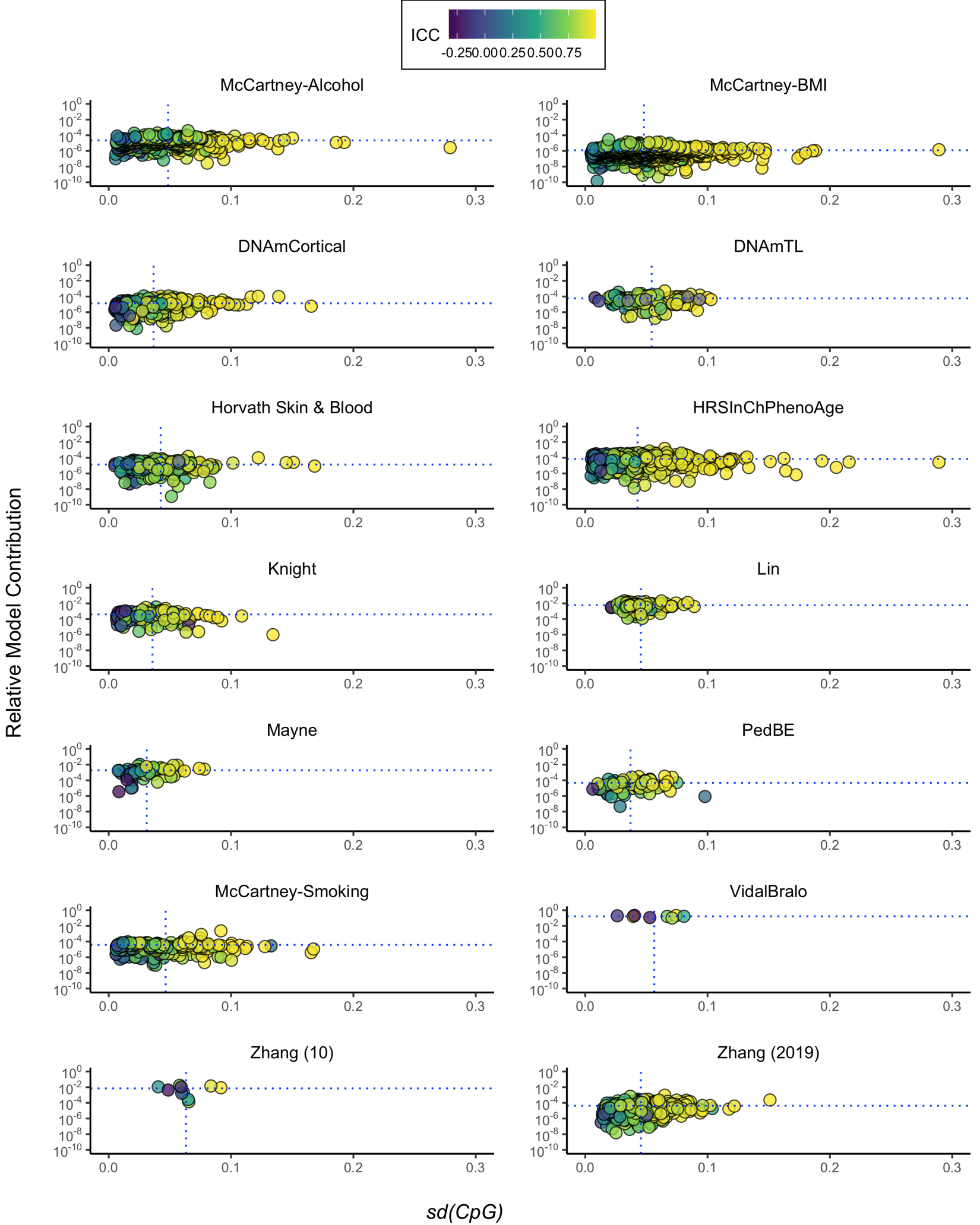

**Figure S1: Extended Comparison of Clock CpGs’ Variability by Model Contribution.** Additional clock examples available within the methylCIPHER package as an extension of **Figure 3A-C**. Individual clock CpGs’ relative contribution to the model weight is plotted against their standard deviation in an FHS offspring cohort, and colored by Beta-value ICC in a technical replicate dataset.

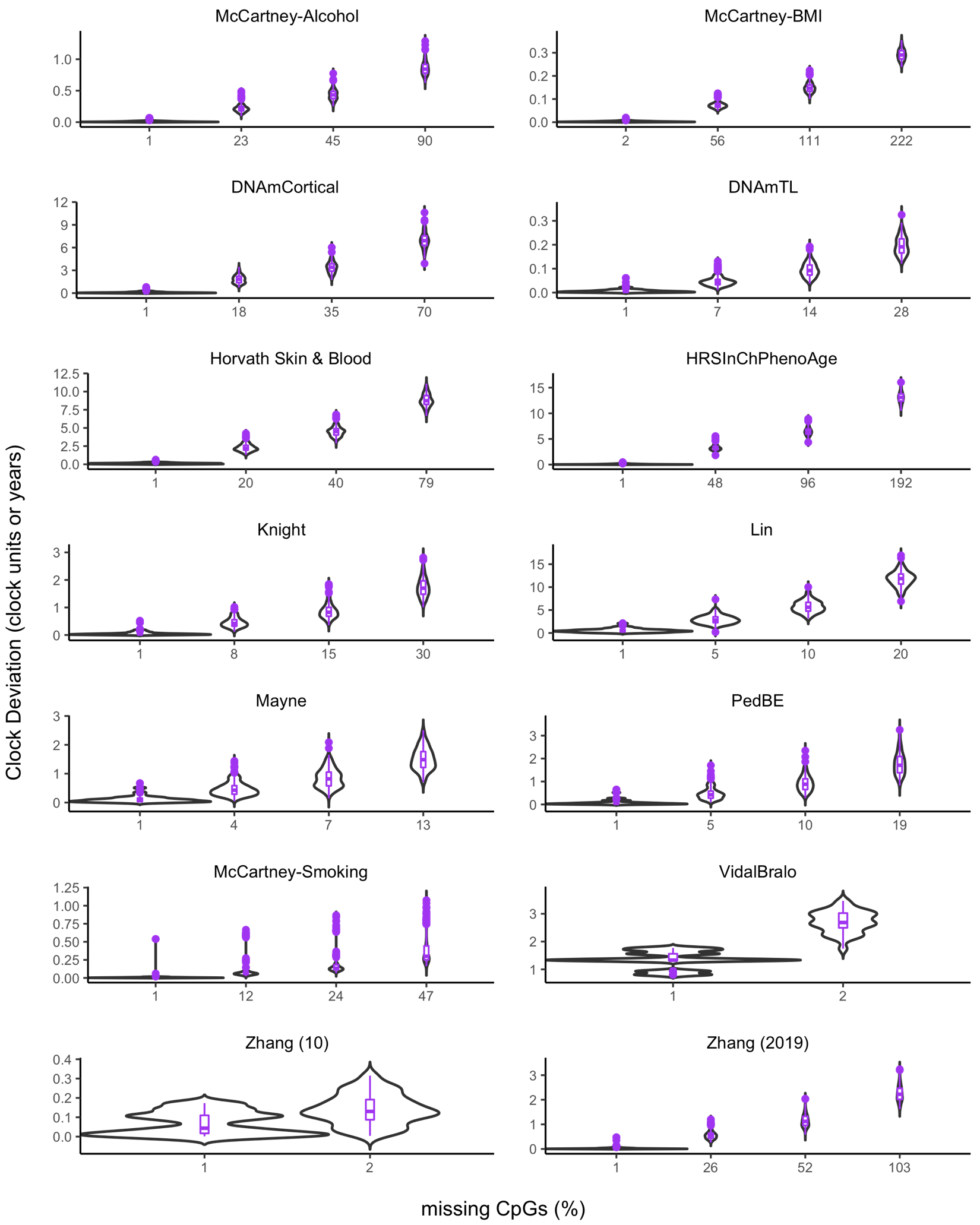

**Figure S2: Extended Comparison of Missed Information in Clock Units According to Increasing Proportions of Missing Data.** Additional clock examples available within the methylCIPHER package as an extension of **Figure 3D-F.** We repeatedly (1000x) tested and plotted the distribution of effects of mean imputation on 0.1%, 5%, 10% and 20% of CpGs selected at random from each clock. Due to the varying sizes of predictors, this represented varying numbers of CpGs, rounded up to the nearest whole CpG. As they only have 10 CpGs, the Zhang and Vidal Bralo clocks’ 0.1%, 5% and 10% trials were all equivalent to just 1 missing CpG. Plots are all visualized as violin plots (black) with quartile boxplots (purple).
